# Membrane potential dynamics of peripheral cold-sensitive neurons in *Drosophila* larvae

**DOI:** 10.64898/2025.12.31.695714

**Authors:** Akira Sakurai, Sergiy M. Korogod, Daniel N. Cox, Gennady S. Cymbalyuk

**Affiliations:** Neuroscience Institute, Georgia State University, Atlanta, GA 30303, USA; School of Life Sciences, Arizona State University, Tempe, AZ 85287, USA

**Keywords:** sensory encoding, thermoception, thermosensation, input resistance, receptor potential

## Abstract

*Drosophila* larvae sense diverse external stimuli through peripheral multidendritic (md) sensory neurons, among which Class III (CIII) neurons are essential for detecting noxious cold temperatures. Previous extracellular recordings showed that CIII neurons produce a characteristic phasic–tonic response to cold: a transient high-frequency spiking phase, during which about half of the neurons burst, followed by frequency adaptation. Here, we report the first intracellular recordings from CIII neurons, allowing us to characterize the membrane potential dynamics underlying their cold-evoked spiking responses. At rest, CIII neurons showed membrane potential fluctuations that occasionally triggered sporadic spikes. Cold stimulation induced a sustained depolarization that evoked a train of spikes, with spike rates peaking at 2–12 spikes/s within the first 5 s, followed by a gradual decline. About half of the neurons exhibited occasional bursts, in which clusters of 2–6 spikes at 10–120 Hz superimposed on a brief depolarizing hump. Cold stimulation also increased input resistance and amplified electrotonically spreading spikes, suggesting enhanced electrotonic signaling from distal sites via an increased space constant. Together, these findings provide key insights into the electrophysiological mechanisms of sensory encoding in peripheral neurons and establish a framework for dissecting the ionic and structural mechanisms of cold transduction.

**New & Noteworthy:** We demonstrate the first intracellular recordings from *Drosophila* Class III primary sensory neurons, revealing how cold stimuli are encoded at the membrane level. Cold temperature evokes sustained depolarization and trains of action potentials, with roughly half of neurons producing bursts. This activity is accompanied by increased input resistance, likely enhancing electrotonic signals from distal loci. Cold encoding thus involves more complex mechanisms than previously recognized, combining dendritic activation with global changes in membrane conductance.

## Introduction

The primary sensory cells serve as the first interface through which animals acquire information from the external environment across diverse sensory modalities. Through signal transduction, these cells convert sensory stimuli into neuronal signals that can be processed by the nervous system (Delcomyn 1998; Dhaka et al. 2006; Julius and Nathans 2012). The stimulus-evoked responses of sensory neurons have been investigated extensively at both the cellular and population levels, providing critical insights into how sensory inputs are represented as patterns of neural activity. Extracellular recordings have revealed stimulus-evoked activity in single or small groups of sensory neurons (Gallar et al. 2003; Kashkoush et al. 2019; Maksymchuk et al. 2022; Song et al. 2007; Yan et al. 2013), while imaging with calcium- and voltage-sensitive dyes enables recording of activity across neuronal populations (Babes et al. 2004; Patel et al. 2022; Ran et al. 2016; Shannonhouse et al. 2025; Terada et al. 2016; Walters et al. 2019; Wang et al. 2018).(Babes et al. 2004; Patel et al. 2022; Ran et al. 2016; Shannonhouse et al. 2025; Terada et al. 2016; Walters et al. 2019; Wang et al. 2018). To understand how sensory input alters the membrane potential of neurons and how these changes are translated into patterns of action potentials, it is essential to directly record and manipulate the membrane-potential responses. In mammals, the dynamic membrane properties of sensory neurons have been first investigated by severing their peripheral axons, which induces ectopic discharge near the somata in the dorsal root ganglia (Amir and Devor 1997; Amir et al. 2002a; Amir et al. 1999; Gossard et al. 1999; Reid and Flonta 2001a). In invertebrate sensory neurons, the somata are located adjacent to the peripheral dendrites, but the surrounding sheath makes direct intracellular recordings particularly challenging (Omoto et al. 2016; von Hilchen et al. 2013). Consequently, only a few studies have succeeded in directly recording membrane potentials from peripheral sensory neurons (Cao et al. 2016; Hardie 1991a; Laurent et al. 2002; Ripley et al. 1968; Seyfarth and French 1994).

In *Drosophila* larvae, peripheral multidendritic sensory neurons play a crucial role in mediating modality-specific behavioral responses to external stimuli (Tracey et al., 2003; Zhong et al., 2012; Ohyama et al., 2013; Terada et al., 2016; Yoshino et al., 2017). Among them, class III (CIII) neurons are primarily responsible for sensing cold and innocuous mechanical stimuli (Maksymchuk et al. 2022; Patel et al. 2022; Tsubouchi et al. 2012; Turner et al. 2016; Yan et al. 2013), whereas class IV (CIV) neurons are responsible for triggering nociceptive responses to heat, chemical, and noxious mechanical stimuli (Himmel et al. 2019; Lopez-Bellido et al. 2019; Ohyama et al. 2013; Terada et al. 2016; Tracey et al. 2003; Yoshino et al. 2017; Zhong et al. 2012). Their firing patterns and molecular mechanisms have been characterized using experiments that combine gene knockdown approaches with calcium imaging and extracellular recordings of action potentials (Himmel et al. 2019; Maksymchuk et al. 2022; Maksymchuk et al. 2023; Onodera et al. 2017; Patel et al. 2022; Terada et al. 2016; Turner et al. 2016; Yan et al. 2013). In CIII neurons, transient receptor potential (TRP) channels, SLC12 family chloride exchangers, calcium-activated chloride channels, metabotropic GABAergic signaling pathways, and calcium-induced calcium release mechanisms have been shown to play important roles (Himmel et al. 2023; Patel et al. 2022; Turner et al. 2016).

Here, we performed direct intracellular recordings of CIII neurons to characterize their membrane potential dynamics. At room temperature, these neurons exhibited membrane potential fluctuations that occasionally triggered action potentials or groups of spikes, *i.e*., bursts. Cold stimulation induced a sustained depolarization accompanied by increased input resistance, resulting in a barrage of action potentials. Notably, nearly half of the neurons generated bursts, which are characterized by multiple high-frequency spikes superimposed on an underlying depolarizing hump. These findings provide new insights into the membrane potential dynamics of CIII neurons and establish a foundation for future studies on the mechanisms of sensory transduction and computational models of sensory encoding of temperatures.

## Materials and Methods

### Animals

*Drosophila melanogaster* larvae were reared on a standard cornmeal-molasses-agar diet under a 12:12-hour light-dark cycle at room temperature (21–24°C) (Himmel et al. 2022). To visualize CIII neurons using fluorescence microscopy, larvae expressing *UAS-mCD8::GFP* (Bloomington *Drosophila* Stock Center #5130, Indiana University, Bloomington, IN) under the control of *GAL4-19-12* were used (**Fig. 1A**). The *GAL4-19-12* line was kindly provided by Dr. Y.-N. Jan (Xiang et al. 2010).

**Figure 1.**
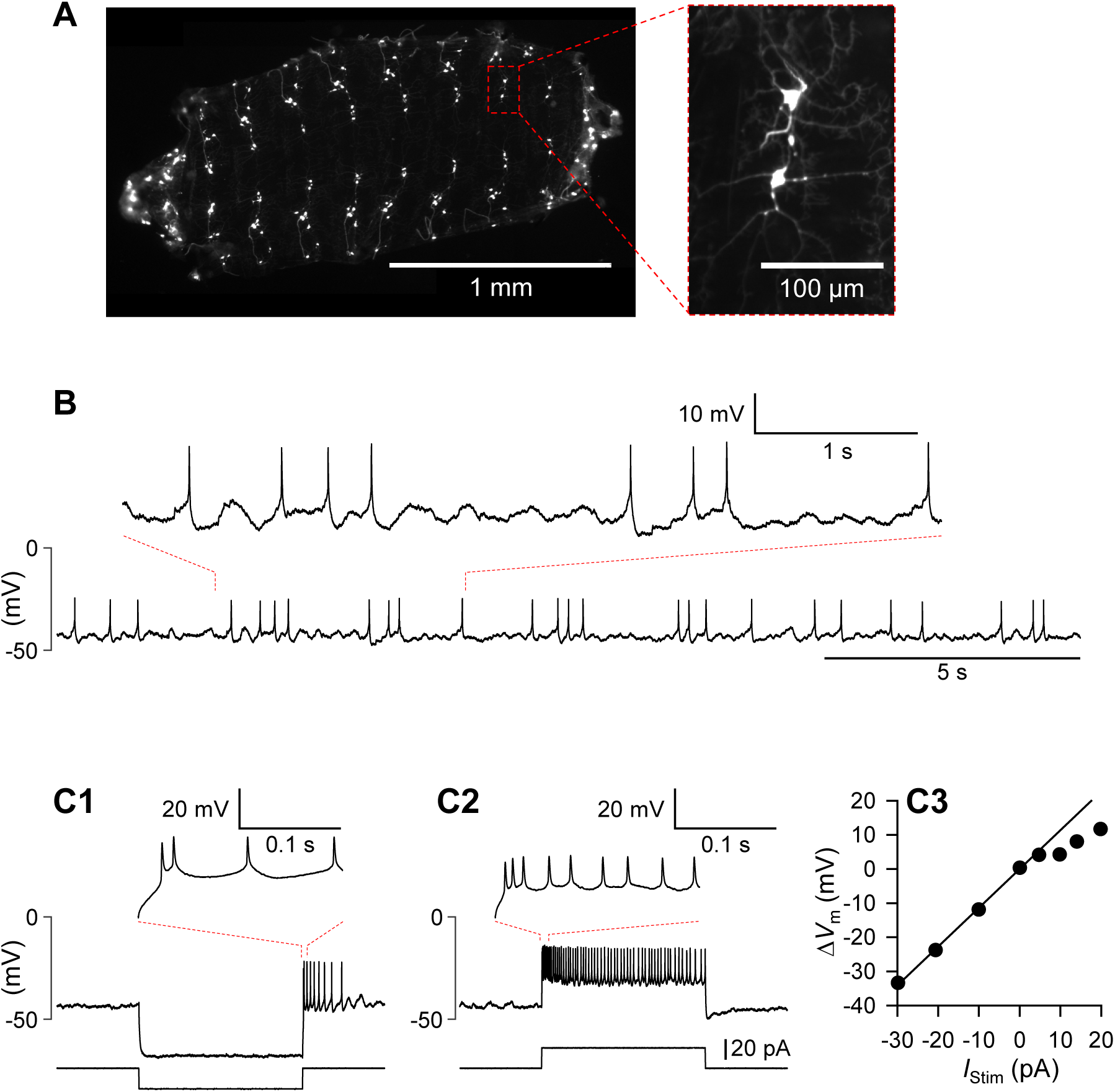
Spontaneous membrane potential activity in a CIII neuron. ***A***, Fluorescence micrograph of a fillet preparation of a *Drosophila* larva, showing GFP-expressing CIII neurons. ***B***, CIII neurons exhibited irregular fluctuations in membrane potential, with action potentials riding on depolarizing fluctuations. ***C1-3,*** Voltage responses of a CIII neuron to a hyperpolarizing (***C1***) and a depolarizing (***C2***) current pulse, and an I–V relationship of the same neuron, plotting injected current versus resulting membrane potential change (***C3***). The input resistance of the neuron was calculated from the slope of the linear portion of the I–V relationship.

### Electrophysiology

Second instar larvae, 2–4 mm in body length, were dissected as described previously (Himmel et al. 2022; Maksymchuk et al. 2022). Each larva was immobilized in a Sylgard-lined dish (Sylgard 184, Dow Corning, Midland, MI, USA) by pinning the proboscis and tail. The ventral body wall was then cut longitudinally and pinned flat with the inner side facing upward. The dissecting dish was transferred to the stage of a fluorescent microscope (Axioplan, Carl Zeiss, Oberkochen, Germany), and the somata of CIII neurons were viewed through a water-immersion lens (W N-Achroplan 63×/0.9 M27, Carl Zeiss, Oberkochen, Germany). Protease (0.5%; type XIII, Sigma, St. Louis, MO, USA) dissolved in saline was applied by gentle pressure ejection from a pipette with a tip diameter of approximately 20 µm to remove the muscle while the preparation was continuously superfused with saline. Throughout the experiment, the dish was continuously superfused with gravity-fed saline composed of (in mM): NaCl 120, KCl 3, MgCl₂ 4, CaCl₂ 1.5, NaHCO₃ 10, trehalose 10, glucose 10, sucrose 10, TES 10, and HEPES 10. The saline osmolality was 305 mOsm, and pH was adjusted to 7.2.

CIII neurons were identified using epifluorescence microscopy by detecting GFP fluorescence, which was driven by *GAL4-19-12*-mediated expression of *UAS-mCD8-GFP.* For the intracellular recordings, pipettes were pulled from borosilicate glass tubes with filaments (OD: 1 mm, ID: 0.5 mm) using an electrode puller (P-97, Sutter instrument, Novato, California, USA), resulting in the resistances of 30–60 MΩ when filled with the electrode solution. The composition of the electrode solution was as follows (in mM): 185 K-gluconate, 5 NaCl, 2 MgCl₂, 0.1 CaCl₂, 1 EGTA, and 10 HEPES (pH 7.4, ∼390 mOsm). The electrodes were connected to the amplifier headstage (Gain x0.1LU; AxoClamp 2B, Molecular Devices, San Jose, CA, USA) via Ag-AgCl wire. To penetrate the cell membrane, the electrode tip was first positioned against the cell, followed by the application of suction using a 10-mL syringe. When the tip resistance exceeded several GΩ, negative current pulses (100–1000 pA, 1–50 ms) were applied to rupture the membrane. Amplifier output signals were digitized at 10 kHz using an A/D converter (Micro1401-4, Cambridge Electronic Design, Cambridge, UK) and recorded on a PC running Spike2 software (version 8, Cambridge Electronic Design, Cambridge, UK) under Windows 10 (Microsoft, Redmond, WA, USA).

To apply cold temperature stimulation, the perfusion was switched to saline cooled to the desired temperature, delivered directly onto the preparation from the outlet of an in-line solution cooler (SC-20, Warner Instruments, Hamden, CT, USA) positioned immediately upstream. The cooler was set in advance to the target temperature (10°C) to chill the saline before delivery. This protocol allowed the local temperature in the vicinity of the preparation to decrease at a rate of -2 to -6°C/sec. Saline temperature was continuously monitored using a BAT-12 microprobe thermometer (Physitemp, Clifton, NJ, USA), with the temperature probe positioned adjacent to the fillet preparation. Temperature readings were simultaneously transmitted to the Micro1401. In some experiments, tetrodotoxin (TTX; Sigma-Aldrich, St. Louis, MO) was used to abolish spikes; it was prepared as a 1 mM stock solution in distilled water, aliquoted, stored at −20°C, and diluted to 1 μM immediately before use.

To measure the somatic input resistance of CIII neurons, AxoClamp 2B was set to the discontinuous current clamp mode, and 4-second hyperpolarizing current pulses of different amplitudes (−30, −20, −10 pA) were injected through the recording electrode. The resulting membrane potential changes were measured, and input resistance was calculated from the slope of the regression line obtained by plotting voltage amplitude against injected current (**Fig. 1C3**). To monitor changes in input resistance during cold stimulation, brief current pulses (0.2 or 0.5 s, −10 or −20 pA) were applied once per second before and throughout the stimulus. Percent changes in input resistance were calculated from the average of five voltage deflections elicited by current pulses before and near the end of cold stimulation.

### Data analysis and statistics

To detect spikes in the recorded data, we set the amplitude threshold using the Spike2 software function. Spiking rate (spikes per second) was calculated as the average over a fixed time window (1 s). Statistical comparisons were performed using SigmaPlot version 12.5 (Jandel Scientific, San Rafael, CA) for two-tailed paired t-tests. Results are presented as the mean ± standard deviation.

## Results

### Membrane potential activity of CIII neurons

CIII neurons exhibited a resting membrane potential ranging from –53 mV to –29 mV, with a mean of –41.9 ± 5.3 mV (mean ± SD, N = 46). The membrane potential was not stable but consistently showed irregular fluctuations with amplitudes below 4 mV, averaging 2.3 ± 0.41 mV (mean ± SD, N = 46; **Fig. 1B**). Action potentials occurred when these fluctuating depolarizations reached threshold. The lack of overshooting spikes suggests limited regenerative spike mechanisms in the somatic membrane, consistent with action potentials being initiated in more distal compartments and spreading passively to the soma.

To assess the membrane properties of CIII neurons at room temperature, square current pulses of varying amplitudes were injected (**Fig. 1C1, C2**). Spikes were evoked upon release from a 4-s hyperpolarization induced by a negative current pulse (**Fig. 1C1**). This rebound excitation often produced spike doublets with short interspike intervals from 0.005 to 0.124 s (-30 pA pulse, 0.017 ± 0.016 s, N = 29 of 44; -20 pA pulse, 0.017 ± 0.018 s, N = 25 of 46; -10 pA pulse, 0.039 ± 0.037 s, N = 11 of 44; all values are mean ± SD; **Fig. 1C1**, inset). Depolarizing current pulses elicited sustained trains of action potentials, with a high initial spiking rate that gradually declined. The minimal interspike intervals ranged from 0.006 to 0.079 s (10 pA pulse, 0.029±0.016 s, N = 45; 20 pA pulse, 0.014±0.008 s, N = 42; all values are mean±SD). From the I–V relationship obtained with hyperpolarizing current pulses (**Fig. 1C3**), the input resistance was 2.10 ± 0.96 GΩ (mean ± SD, N = 21).

### Membrane potential responses to cold temperature

In previous studies (Maksymchuk et al., 2022, 2023), cold stimulation was shown to elicit prominent peaks of spiking activity in CIII neurons. Here, we observed that the cold-evoked spiking occurred on top of an underlying slow depolarization (**Fig. 2A1, A2**).

**Figure 2.**
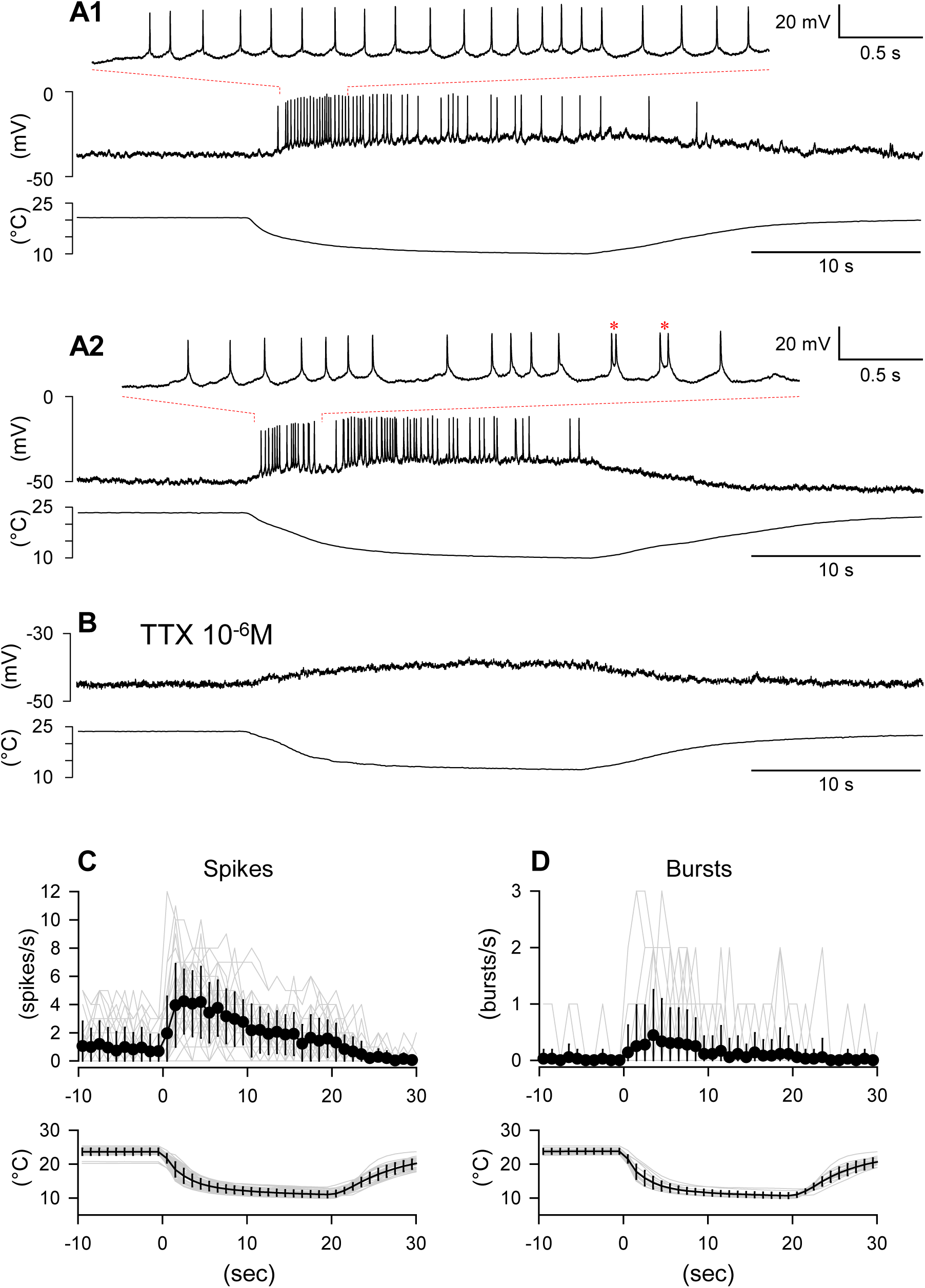
Membrane potential responses of CIII neurons to cold stimulation. ***A1*-*2***, Simultaneous recordings of the membrane potential of a CIII neuron (top) and the perfusate temperature (bottom). In ***A1***, a train of tonic spikes was observed, whereas in ***A2***, occasional bursts of multiple spikes riding on single depolarizing humps (red asterisks) occurred among the tonic spikes. For both ***A1*** and ***A2***, an expanded trace of the region outlined by red dashed lines is shown as an inset above. ***B***, Application of TTX abolished the fast spikes, leaving only a gradual depolarizing response throughout the stimulation. ***C***, Spike rate and temperature plotted in 1-s bins. Mean spike rates (upper panel) and temperature (lower panel) are shown as black circles and a line, respectively. Error bars indicate standard deviations. ***D***, Burst rate and temperature plotted as in ***C***.

In 55.6% of CIII neurons (N=20 of 36), a 20-second cold stimulation induced tonic spiking responses (**Fig. 2A1**). The remaining 45.4% (N=16 of 36) exhibited occasional bursts, where two or more spikes were superimposed on a brief depolarizing hump during the tonic spiking activity (**Fig. 2A2**). Application of TTX (10⁻⁶ M) eliminated all the spikes, but the gradual depolarization was still seen during the stimulus (**Fig. 2B**). Spike rate, on average, reached the maximum within five seconds after stimulus onset and then gradually declined (**Fig. 2C1**). For tonic spikes, minimal interspike intervals ranged from 0.055 to 0.59 s (0.16 ± 0.11 s, N = 36), and maximal intervals from 0.31 to 6.98 s (2.16 ± 1.50 s, mean ± SD, N = 36). In bursting neurons, intra-burst interspike intervals ranged from 0.008 to 0.11 s (0.040 ± 0.015 s, mean ± SD, N = 16). On average, the spike rate during the first half of the stimulation (0–10 s; 3.44 ± 1.69 spikes/s) was significantly higher than during the latter half (10–20 s; 1.80 ± 1.29 spikes/s; mean ± SD, N = 36; two-tailed paired t-test, P < 0.001). The rate of burst occurrence was also higher during the first 10 s (0.61 ± 0.56 bursts/s) than the latter 10 s (0.23 ± 0.21 bursts/s; mean ± SD, P = 0.012 by two-tailed paired t-test, N =16; **Fig. 2D**). The average intra-burst spike rate during the first 10 s was 50.59 ± 30.18 spikes/s (N = 14), which decreased significantly to 33.18 ± 22.69 spikes/s during the latter 10 s (N = 13), a difference that was also significant (P = 0.002 by two-tailed paired t-test).

### Changes in membrane input resistance during cold stimulation

Turner et al. (2016) implicated select TRP channels in mediating cold-evoked behavior in Drosophila larvae, suggesting these channels contribute to the cold-evoked excitatory response of CIII neurons. To examine changes in membrane properties during cold-evoked depolarization, we manipulated the membrane potential with constant-current injections of varying amplitudes and observed the effects on the depolarizing response (**Figs. 3A1–A3**).

**Figure 3.**
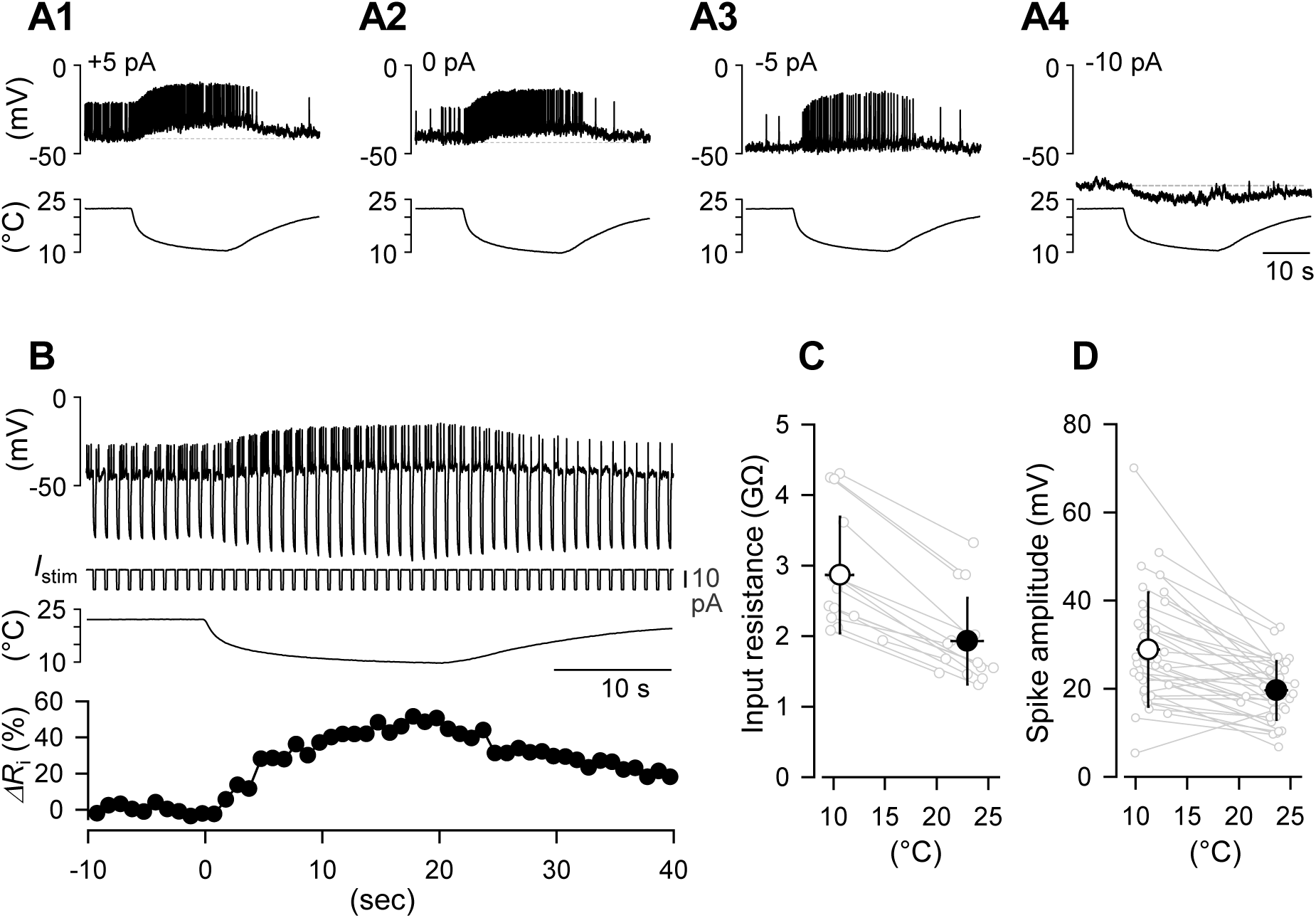
Membrane potential responses to cold stimulation and changes in input resistance. ***A1-4***, Responses to cold stimulation at different hyperpolarizing current injections: ***A1***, +5 pA; ***A2***, control; ***A3***, –5 pA; ***A4***, –10 pA. ***B***, Changes in input resistance of CIII during cold stimulation. From top to bottom: membrane potential response to repeated hyperpolarizing current pulses (-20 pA, 0.2 s) applied at 1 Hz (*I*_stim_), change in temperature, and the corresponding plot of % changes in input resistance. ***C***, Plots of input resistances at room temperature (black circle) and during cold stimulation (open circle). Individual data were shown as gray symbols. Error bars indicate standard deviations. ***D***, Spike amplitude at two different temperatures plotted as in ***C***.

The amplitude and polarity of the cold-evoked response at the soma depended on the membrane potential set by the injected current. Hyperpolarization reduced spike rate and suppressed the slow depolarization (**Fig. 3A3**), and stronger hyperpolarization inverted the response, producing a hyperpolarizing potential during cold stimulation (**Fig. 3A4**). In contrast, depolarizing current injection enhanced the slow depolarization (**Fig. 3A1**). To further probe changes in membrane properties, we applied repetitive negative current pulses (10 pA for 0.2 s) to monitor input resistance (**Fig. 3B**). During cold stimulation, the voltage change induced by these pulses increased in amplitude, indicating a sustained increase in input resistance. On average, input resistance increased significantly from 1.93 ± 0.61 GΩ to 2.94 ± 0.82 GΩ during cold stimulation (P = 0.0003 by two-tailed paired t-test, N = 15; **Fig. 3C**). The amplitude of the spikes also increased during cold stimulation from 19.63 ± 6.66 mV to 28.89 ± 12.99 mV (P < 0.001 by two-tailed paired t-test, N = 35; **Fig. 3D**). These results suggest that the somatic slow depolarization cannot be explained solely by the opening of nonselective cation channels such as TRP channels; however, it would be consistent with TRP activation if these channels are located at electrically remote dendritic compartments. The sustained increase in somatic input resistance indicates that cold stimulation reduces membrane conductance at least in the soma vicinity. Together, reduced somatic conductance and electrotonic spread from distal transduction sites likely contribute to the composite cold-evoked response, with the increase in input resistance enhancing the spread of depolarization to the soma.

## Discussion

In this study, we directly recorded the membrane potential of CIII neurons, a type of multidendritic primary sensory neuron located beneath the body wall of *Drosophila* larvae, using sharp intracellular electrodes to observe their responses to cold stimulation. While direct recordings of membrane potential activity have been reported in other arthropod sensory neurons, such as isolated insect photoreceptor cells, (Hardie 1991b; Juusola and Hardie 2001), mechanoreceptor neurons in cockroach sensory hairs (Stockbridge and French 1991), mechanoreceptors in spider legs (Seyfarth and French 1994), and olfactory receptor cells in honeybee (Laurent et al. 2002) and fly (Cao et al. 2016), to our knowledge this is the first study to directly record the membrane potential responses of the primary sensory neurons to temperature changes.

At rest, the soma of CIII neuron exhibited subthreshold membrane potential fluctuations of varying amplitudes, occasionally triggering spontaneous discharges. These voltage fluctuations likely underlie the irregular spikes observed in extracellular recordings from previous studies (Maksymchuk et al. 2022). Similar subthreshold oscillations and clustered spikes have been reported in mammalian dorsal root ganglion neurons (Amir and Devor 1997; Amir et al. 2002a; Amir et al. 2002b; 1999; Bade et al. 1979; Braun et al. 1980; Schafer et al. 1982) and trigeminal ganglion neurons (Wu et al. 2001). In these mammalian neurons, oscillations arise from the interplay between persistent Na⁺ channels and voltage-gated K⁺ currents, with TTX-sensitive Na⁺ spikes riding on top of the membrane potential oscillations (Amir et al. 2002a; Amir et al. 1999; Wu et al. 2001). Additionally, a slow depolarizing afterpotential (DAP) following hyperpolarization contributes to the generation of spike trains or bursts (Amir et al. 2002b). The mechanisms underlying membrane potential fluctuations in CIII neurons remain unknown and are beyond the scope of this study. The rebound discharges observed upon release from hyperpolarizing current injections (**Fig. 1C1**) may indicate the involvement of hyperpolarization-activated currents or currents de-inactivated by hyperpolarization, such as hyperpolarization-activated cyclic nucleotide–gated (HCN) channels (Mishra and Narayanan 2025) and T- or R-type Ca^2+^ channels (Engbers et al. 2011; Metz et al. 2005).

### Cold-evoked somatic depolarization with increased input resistance

Cold temperature stimulation induced a barrage of action potentials riding on a slow depolarization in CIII neurons. This depolarization was accompanied by an increase in membrane input resistance and appeared to diminish or reverse when the neuron was hyperpolarized (**Fig. 3A2-A4, B, C**). Similar depolarizing responses accompanied by increased membrane resistance upon cooling have been reported across a wide range of species, from unicellular organisms to mammalian neurons (Alamri et al. 2018; Cabanes et al. 2003; Inoue and Nakaoka 1990; Klee et al. 1974; Lee et al. 2005; Viana et al. 2002; Volgushev et al. 2000). These effects are attributed, in part, to the closure of background potassium channels (Maingret et al. 2000; Reid and Flonta 2001a; Reid and Flonta 2001b; Viana et al. 2002). Temperature also strongly influences the kinetics of voltage-gated channels, thereby altering the shape of action potentials (Frankenhaeuser and Moore 1963; Hodgkin and Huxley 1952; Joyner 1981; Klee et al. 1974; Sarria et al. 2012; Thompson et al. 1986). In mammalian sensory afferents, K2P channels mediate cooling-induced reductions in leak conductance, increases in input resistance, membrane depolarization, and changes in action potential waveform and timing (Kanda et al. 2021). In this study, the apparent reversal of the slow depolarization during hyperpolarization may predominantly reflect a passive response to the injected hyperpolarizing current due to the increase in input resistance. The closure of potassium channels likely contributes to the cold-evoked depolarization; however, contributions from other cation or chloride channels with different temperature dependencies, which may shift the balance of inward and outward currents, cannot be excluded.

There is growing evidence that cooling can directly activate specialized temperature-sensitive channels, such as those of the TRP family, which open at low temperatures and increase excitability through depolarization (McKemy 2007; McKemy et al. 2002; Okazawa et al. 2002; Peier et al. 2002; Reid and Flonta 2001b; Story et al. 2003). Similarly, studies on temperature sensitivity of *Drosophila* larvae have mainly focused on the involvement of TRP channels (Gallio et al. 2011; Luo et al. 2017; Montell 2021; Sokabe 2025; Turner et al. 2016). In CIII neurons, at least three TRP channels, which are NompC, Pkd2, and Trpm, have been implicated in cold nociception (Turner et al. 2016); although their precise subcellular localization remains unknown.

The increase in membrane resistance during cold stimulation may have two opposing effects. On one hand, it could counteract the additional conductance produced by dendritic receptor channel openings, making it difficult to isolate the receptor potential when recorded from the soma. On the other hand, it increases the electrotonic space constant, thereby reducing attenuation of dendritically generated signals en route to the recording site at the soma. Given the small size of action potentials recorded at the soma, spike initiation likely occurs elsewhere. Consistent with this interpretation, our attempts to measure receptor currents using single-electrode voltage-clamp recordings were unsuccessful (data not shown). This likely reflects the extensive dendritic arborization of CIII neurons, which places the thermoreceptor site electrotonically far from the soma where recordings were made. In the nociceptive CIV neuron, fast sodium channels are predominantly localized in the axon and peripheral dendrites (Rey et al. 2023), and CIII neurons presumably share a similar distribution pattern. The interplay among temperature-dependent changes in membrane conductance, the spatial distribution of these channels, and neuronal morphology likely determines how cold stimuli are encoded into spiking activity. Further experiments that combine calcium imaging with somatic voltage recordings will be needed to clarify how signals are conducted from dendrites to the soma.

### Diversity of activity patterns

Consistent with previous studies (Maksymchuk et al. 2022; Maksymchuk et al. 2023), the spike frequency of CIII neurons peaked shortly after a rapid temperature drop, and then declined during the sustained low temperature. Two distinct types of firing were observed. Nearly half of CIII neurons exhibited bursting activity, in which multiple spikes rode on a slow depolarization, in addition to tonic firing during cold stimulation. The remaining neurons fired only tonic spikes at relatively stable frequencies. This heterogeneity in firing patterns is consistent with our previous findings (Maksymchuk et al. 2023). Bursting has been widely recognized as an important feature in sensory systems of both vertebrates and invertebrates (Krahe and Gabbiani 2004). First, high-frequency trains of spikes within bursts increase the reliability of synaptic transmission by rapidly elevating presynaptic calcium, thereby facilitating temporal summation and boosting neurotransmitter release (Del Castillo and Katz 1954; Lisman 1997; Zucker 1999). Second, bursting activity may improve the signal-to-noise ratio of sensory information and enhance the contrast of transmitted signals (Eggermont and Smith 1996). Together, these mechanisms suggest that bursting plays a key role in optimizing sensory encoding and transmission.

The cellular basis of this diversity in activity patterns remains unclear, but differences in membrane excitability, ionic contributions to membrane potential changes, and cell-to-cell variability are all likely factors. Previous computational modeling combined with extracellular recordings predicted that CIII neurons possess ion channels capable of generating both tonic spiking and bursting even in the absence of thermoTRP activation (Maksymchuk et al. 2023). Based on these models, a critical role was hypothesized for an inward current, likely mediated by Ca²⁺ channels, whose activation and inactivation kinetics drive bursting dynamics. Notably, the square-wave burst waveform predicted by the model closely resembles the bursting activity observed in the present intracellular recordings. Differences in the kinetic properties, expression levels, and subcellular distribution of thermoTRP and voltage-gated channels may therefore underlie whether individual CIII neurons respond to cold with tonic spiking or bursting (Maksymchuk et al. 2022; Maksymchuk et al. 2023). Dendritic morphology may also play an important role, as computational models showed that branching patterns strongly influence neuronal firing dynamics (Korogod and Kaspirzhny 2008; Korogod and Tyč-Dumont 2009; Mainen and Sejnowski 1996).

Taken together, the present results highlight the complex membrane potential dynamics of CIII neurons and the diversity of their activity patterns, advancing our understanding of how peripheral sensory neurons encode thermal stimuli. They also provide a foundation for future studies to test how the composition of thermosensitive TRP channels, channel kinetics, and dendritic architecture determine whether a CIII neuron generates tonic spiking or bursting activity by combining cell-specific knockdowns with computational modeling.

## Conflict of interest

The authors declare no competing financial interests.

## Acknowledgement

This work was supported by NIH grant 2R01NS115209-06 to Daniel Cox and Gennady Cymbalyuk.

## Notes

### Competing Interest Statement

The authors have declared no competing interest.

